# Stimulating prefrontal cortex facilitates training transfer by increasing representational overlap

**DOI:** 10.1101/2023.11.23.568508

**Authors:** Yohan Wards, Shane E. Ehrhardt, Hannah L. Filmer, Jason B. Mattingley, Kelly G. Garner, Paul E. Dux

## Abstract

Difficulties in multitasking may be the price humans pay for our ability to generalise learning to new tasks. Mitigating these costs through training has been associated with reduced overlap of constituent task representations within a task-related brain network. Transcranial direct current stimulation (tDCS), which can modulate neural activity, has shown promise in generalising training gains. Whether tDCS influences the changes in task-associated representations to produce such training generalisation remains unexplored. Here, we paired prefrontal cortex tDCS with multitasking training, and collected functional magnetic resonance imaging data pre- and post-training. We found that 1mA tDCS enhanced visual search performance, and using machine learning to assess the overlap of brain activity related to the training, show that these generalised gains were predicted by changes in classification accuracy for patterns of frontal, parietal, and cerebellar activity in participants who received left prefrontal cortex stimulation. Thus, prefrontal cortex tDCS interacts with training related changes in task representations, potentially driving the generalisation of learning.

## Introduction

Humans routinely show an ability to rapidly adapt to changes in the environment. Such behaviour requires learning generalisation - the transfer of skills from one task to another - which has been the subject of psychological research for over a century^1^. Despite our competency in flexibly applying our cognition to a wide variety of tasks in everyday life, we demonstrate substantial decrements whenever we perform multiple tasks simultaneously, even when they are simple^2-4^. A fundamental trade-off has recently been proposed whereby multitasking costs reflect mechanisms associated with information sharing across novel tasks^5,6^. This trade-off gives rise to an interesting dilemma as improved capacity for both multitasking and generalisable learning abilities are increasingly desirable in the modern world^7^.

Training interventions are thought to mitigate multitasking costs by reducing representational overlap within frontoparietal-subcortical (FP-SC) brain regions that subserve the constituent tasks^8-13^. Transfer is a core aim of many brain training interventions in commercial settings^14,15^, but evidence for the generalisation of purely training-induced improvements to untrained tasks is mixed^16-19^. An approach that has shown promise in enhancing transfer involves combining non-invasive brain stimulation, such as transcranial direct current stimulation (tDCS), with behavioural training interventions^20-22^.

tDCS, which generates an exogenous electric field that alters the activity of underlying neurons^23,24^ and glial cells^25^, has shown promise in enhancing performance on untrained tasks when paired with training on working memory^20,26,27^, cognitive flexibility^28^, mathematics^29^, and multitasking^21,22^. In a related study, Filmer et al^30^ found that anodal stimulation applied to the left prefrontal cortex during learning resulted in faster evidence accumulation for both the training task and visual search transfer task. This suggests that the application of tDCS during cognitive training can improve information processing efficiency on more than just the trained task. Given recent theoretical and computational advances in understanding the interplay between mechanisms underlying multitasking costs and learning generalisation^6,6,31^, combining prefrontal cortex tDCS with multitasking training provides a unique avenue for exploring the neural substrates of transfer.

Here, using multivoxel pattern analysis of functional magnetic resonance imaging (fMRI) data, we sought to investigate how changes in informational overlap for a trained task may underlie tDCS-induced transfer to a spatial attention task. Specifically, we assessed whether the performance transfer elicited by combined training and tDCS can be attributed to greater overlap in task representations, facilitating the sharing of information across tasks. To this end, we used ultra-high field (7T) fMRI to examine how patterns of functional brain activity are influenced by the effects of combined multitasking training and tDCS. Capitalising on the individual variability commonly observed in both behavioural effects of tDCS^32,33^ and unique patterns of neural activity during task^34,35^, we examined the extent to which changes in task representations associated with training and brain stimulation underpin performance transfer. To anticipate, we found that prefrontal cortex tDCS combined with multitasking training induced transfer to an untrained visual search task. Among participants who received left prefrontal cortex tDCS, those who showed a decrease in fMRI decoding accuracy in superior parietal, orbitofrontal, and cerebellar regions after multitasking training also showed better performance on the visual search task. These findings suggest that the overlapping nature of task representations during multitasking may mediate performance transfer, and therefore may reflect a neural substrate of generalisable learning.

## Results

### Overview

We trained 178 participants (aged 18-40; M = 22.86, SD = 3.93 years, 119 females) on a multitasking protocol shown to result in transfer of performance gains to visual search when combined with tDCS^21^. Our pre-registered study (https://tinyurl.com/5h8u72j5) spanned 10 sessions: an initial behavioural assessment, two MRI sessions pre-training, four days of tDCS-assisted training, a post-training MRI session and behavioural analysis, concluding with a follow-up behavioural assessment 30 days later. Changes in performance, indexed by reaction time (RT) or accuracy differences across time, were assessed on a range of behavioural measures of working memory, attention, response inhibition, and multitasking. We focused our analyses on the fMRI component of the imaging data, applying multivoxel pattern analysis (MVPA) to the blood oxygen level dependent (BOLD) signal during single-task performance. Decoding accuracy served as a proxy for the extent of overlap of task-relevant activity, which we examined pre- and post-training. In addition, we adopted an individual differences approach which allowed us to capitalise on the interindividual variability observed in the neurophysiology underlying cognitive task performance^10,35^, as well as training and tDCS outcomes ^36-38^.

Participants were divided into five groups to distinguish between effects of stimulation region, stimulation intensity, and task specificity, as follows:

1. Sham Group: Underwent sham stimulation with multitasking training, serving as the tDCS control.
2. 1 mA LH Group: Received 1 mA stimulation to the left PFC with multitasking training.
3. 1 mA RH Group: 1 mA stimulation to the right PFC with multitasking training, for comparison with the 1 mA LH Group to assess stimulation location specificity.
4. RSVP Group: 1 mA stimulation to the left PFC but trained on a rapid serial visual presentation (RSVP) paradigm, for comparison with the 1 mA LH Group to determine training task specificity.
5. 2 mA LH Group: 2 mA stimulation to the left PFC with multitasking training, to be compared with the 1 mA LH Group for examining effects of different stimulation intensities.

Comparisons were also made between 1 mA LH, 1 mA RH, and 2 mA LH against the sham group for a complete understanding of stimulation effects on both trained and untrained tasks. Bayesian t-tests were used for all planned comparisons, with Bayes Factor (BF) values indicating the strength of the evidence for the alternate hypothesis (BF_10_ > 3.2)^39^. Only Bayesian results are reported (instead of both Bayesian and NHST tests, as originally preregistered), as they are fundamentally more conservative and so reduce the likelihood of type 1 errors^40^. Nevertheless, the pattern of results using both Bayesian and NHST methods were consistent.

### Combined tDCS and multitasking training led to transfer

Full details of analyses and results, which are beyond the scope of the present investigation, can be found in Wards et al^41^. Here we recap the relevant behavioural findings. We first ensured comparable baseline performance across the participant groups via a pseudo-randomisation procedure for participant allocation. Our analyses revealed strong evidence against pre-training differences among the groups for the relevant trained and untrained task metrics (BF_10_ < 0.085 for all comparisons^41^).

Turning to training effects, multitasking training enhanced performance as indicated by a larger reduction in reaction times pre- to post-training for both single- and multi-task trials for the 1mA LH group compared with the RSVP training group (BF_10_ = 38.182, error < 0.001%; BF_10_ = 313.222, error < 0.001%). There were only small differences in changes in multitasking cost (the difference between reaction times on single stimuli trials compared with reaction times on the dual stimuli trials) between 1mA LH multitasking training vs the RSVP group (BF_10_ = 2.640, error < 0.001%). Notably, despite observing an impact of tDCS on transfer performance (see below), neither tDCS site nor tDCS intensity differentially impacted multitasking performance improvements. Specifically, comparing the 1 mA LH, 1 mA RH, and 2 mA LH groups to sham revealed no effect of stimulation on performance (single-task RTs: all BF_10_ = 0.248 – 0.275, error ≤ 0.009%; multi-task RTs: all BF_10_ = 0.247 – 0.620, error ≤ 0.01%; multitasking cost: all BF_10_ = 0.293 – 1.190, error ≤ 0.009%; see Wards et al^41^).

Crucially, we did observe transfer effects. To investigate the influence of paired training and stimulation on transfer effects, performance on each task for the 1 mA LH, 1 mA RH, and 2 mA LH groups were compared with sham across the entire behavioural battery. Participants receiving 1 mA stimulation to the left or right prefrontal cortex, in conjunction with multitasking training, showed enhanced performance in the visual search task, compared with sham (Figure 1). Specifically, these gains were observed for reaction times on trials with set sizes 12 and 16 (BF_10_ = 11.410, BF_10_ = 13.215 (1 mA LH vs sham); BF_10_ = 14.915, BF_10_ = 122.788 (1 mA RH vs sham), errors all < 0.003%). Furthermore, these effects endured for 1-month post-training for set-size 16 for both the left and right 1 mA prefrontal cortex stimulation groups relative to sham (BF_10_ = 6.080 and 54.284 respectively, errors < 0.004%; see Wards et al^41^).

**Figure 1.**
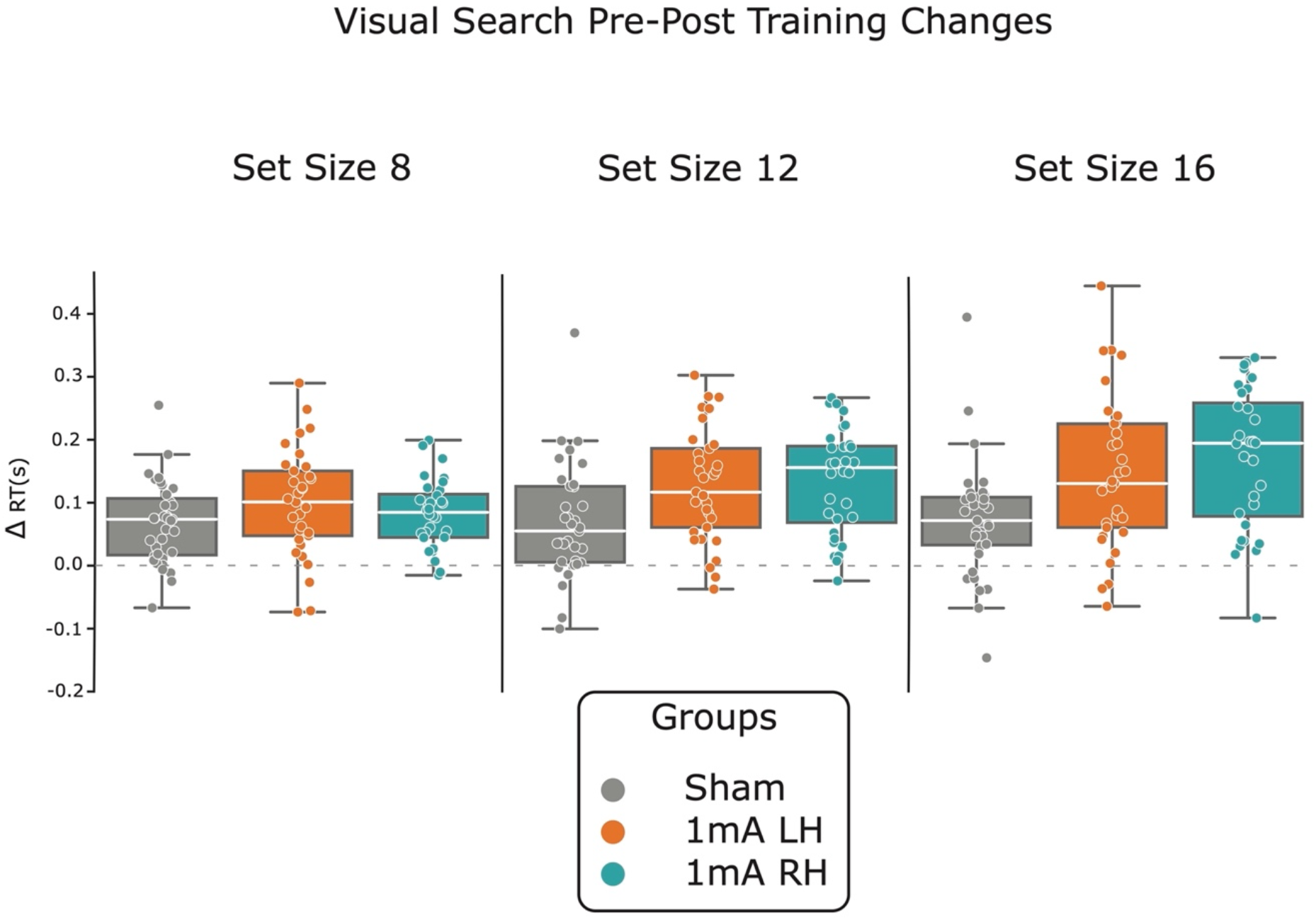
Prefrontal tDCS Results in Enhanced Visual Search Performance. Changes in visual search by group and set size. Sham, and both left and right hemisphere 1mA tDCS groups (1mA LH and 1mA RH respectively) showed similar changes in reaction times for trials with 7 distractors (set size 8). However, both 1mA tDCS groups showed greater performance improvements than sham for set sizes 12 and 16. Each point represents an individual’s change in reaction time from pre- to post-training. Positive values reflect an improvement in performance from pre- to post-training. Δ = change in, RT = reaction time, s = seconds.

### Changes in decoding accuracy underly transfer

Our primary objective was to ascertain whether changes in decoding accuracy within task-relevant multitasking regions predicted changes in performance on the transfer task. Prior research has found that improvements in multitasking via training are associated with a separation of the neural activity patterns corresponding to single-task performance, in a frontal-parietal-subcortical subset of brain regions^10^. Activity within such areas scales with the complexity of many cognitive tasks^42-45^, including visual search^46^. Moreover, Filmer, et al^30^ found that left prefrontal cortex anodal stimulation resulted in faster evidence accumulation for both a response selection task and visual search task. We therefore hypothesised that paired training and tDCS specific alterations in the representation of the training task could underpin generalised performance improvements in visual search.

To isolate task-relevant regions, we tested for brain regions that showed increased activity for both single-tasks from the multitasking training paradigm, relative to fixation (conjunction contrast). This resulted in identification of 41 regions across frontal, parietal, cerebellar, and basal ganglia locations (see S1 Table). These regions broadly align with those identified in earlier studies investigating executive operations^42-45^.

Having identified regions of interest (ROI) that responded to both tasks, we next sought to identify which of those regions showed training-related changes in task representation that corresponded to performance changes on the visual search task. To achieve this, we trained a linear support vector machine algorithm to classify between single-task activity patterns, using voxels in each ROI as features. We then calculated the change in decoding accuracy from pre- to post-training for each ROI and correlated these changes with performance changes on the visual search task within each group. These correlations were compared with sham using the Fisher-Z transformation method, and p-values were corrected for multiple corrections using the false discovery rate (FDR) method.

There was moderate to strong evidence in a subset of the identified regions (Figure 2A), which showed that increases in decoding accuracy pre- to post-training were negatively correlated with improvements in visual search (Figure 2B-F) for individuals who received 1 mA left prefrontal stimulation. Specifically, enhancements in visual search performance corresponded with tDCS- and training-induced decreases in the differentiability of single-task activity in these regions. The anterior cerebellum vermis VI lobe, two left hemisphere superior parietal lobe regions, the right inferior parietal lobe, and the right orbitofrontal cortex all showed negative correlations between changes in decoding accuracy and changes in visual search set size 12 reaction times (r = -0.50, BF_10_ = 11.1; r = -0.45, BF_10_ = 5; r = -0.49, BF_10_ = 10.2; r = -0.45, BF_10_ =5.2; r = -0.58, BF_10_ = 64.7, for each region respectively). Finally, one of the left hemisphere superior parietal lobe regions also showed evidence for a negative correlation between decoding accuracy changes and set size 16 RTs (r = -0.48, BF_10_ = 7.6). Except for the right inferior parietal lobe, all the correlations in the regions presented here were different from sham (FDR corrected p < 0.05 for all comparisons). Notably, no correlations between changes in decoding accuracy and changes in visual search performance were found for the right hemisphere 1mA prefrontal stimulation group.

**Figure 2.**
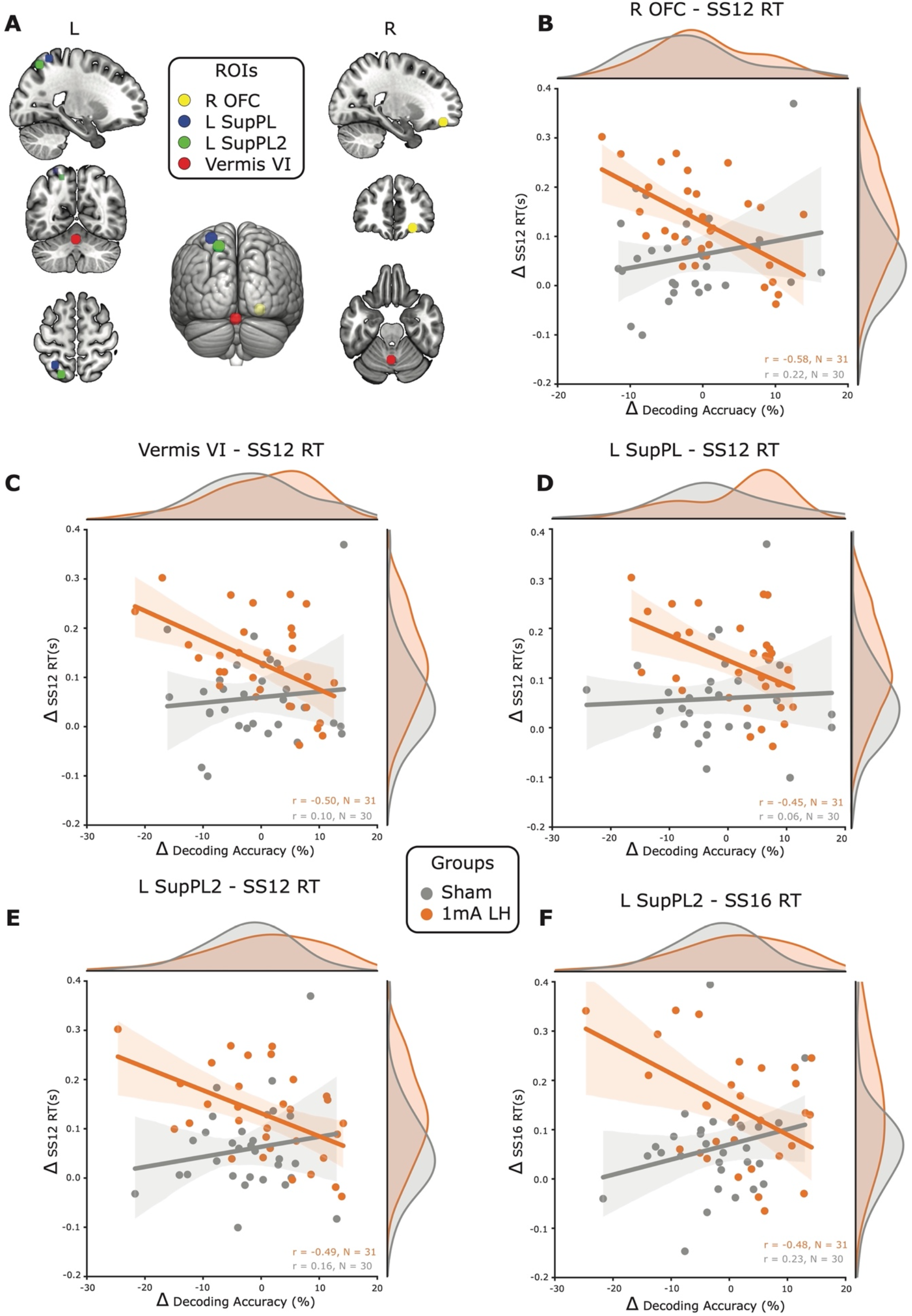
Decreased decoding accuracy for the trained task is associated with improvements on untrained visual search performance. **A**) Regions of interest (ROIs) that were conjointly activated by the trained task subcomponents, and that showed significantly greater correlation coefficients between changes in decoding accuracy and visual search performance changes than sham. These regions were the right orbitofrontal cortex (R OFC - yellow), cerebellum vermis VI (Vermis VI - red), and two regions in the left superior parietal lobe (L SupPL and L SupPL2 – blue and green respectively). **B**) For the group that received left prefrontal cortex anodal tDCS, the R OFC changes in decoding accuracy were negatively correlated with changes in reaction times on visual search trials with 11 distractors (SS12), and this correlation was significantly different from sham. This same pattern of results with negative correlations between changes in decoding accuracy and changes in set size 12 reaction times was observed for the Vermis VI (**C**), L SupPL (**D**), and L SupPL2 (**E**). The L SupPL2 region also had a negative correlation between changes in decoding accuracy and changes in set size 16 reaction times (**F**). Shading around the regression lines displays 95% confidence intervals. L = left, R = right, r = Pearson’s correlation coefficient, n = sample size, s = seconds, Δ = change in.

### Changes in decoding accuracy did not predict multitasking cost improvements

Following from Garner and Dux^10^, we combined all 4 groups that undertook multitasking training and compared them with the group that trained on the RSVP task. As the two groups differed significantly in sample size (MT; n = 122, RSVP; n = 32), we used Levene’s test to check the equality of the variance within each ROI between the two groups (p > 0.05 for each comparison). Next, we averaged the change in decoding accuracy across the frontal-parietal-subcortical (FP-SC) regions in our sample that were the closest (in MNI coordinates) to those identified in Garner and Dux. We then calculated the correlation between changes in multitasking cost and changes in decoding in this FP-SC network within the combined multitasking group (MT), and the RSVP training group. In contrast to Garner and Dux, we found evidence for there being no correlation between changes in decoding in the FP-SC network and changes in multitasking cost for both groups (MT, r = 0.121, BF_10_ = 0.271; RSVP, r = -0.045, BF_10_ = 0.222). This lack of evidence for a change in representational content related to the multitasking component tasks within this FP-SC system in these groups is perhaps not surprising given we did not observe meaningful improvements in multitasking cost for any group that received tDCS. This likely reflects the fact that only approximately one-third of the number of training trials used in Garner and Dux were used here, although there were other design differences, so definitive conclusions cannot be drawn on this issue.

## Discussion

We examined how combined prefrontal tDCS and multitasking training changes neural representations of the trained tasks to give rise to generalisable learning. We found that pre- to post-training decreases in the discernability between trained task representations were associated with faster visual search performance for the group that received left prefrontal tDCS. This association was specific to decoded neural activity in the left superior parietal lobe, right orbitofrontal cortex, and the cerebellar vermis. These results are in line with recent theories of multitasking and learning, which suggest that shared task-representations allow for better knowledge generalisation at the cost of poor multitasking ability^5,6^.

Using a preregistered study design, large sample size (n = 178), double blinding of stimulation conditions, and active control groups, we are able rule out common issues that can limit conclusions from tDCS studies^33,48^. Specifically, we double blinded the 1 mA LH and sham groups, so differences in performance between these two groups cannot be attributed to peripheral causes. Our active training control group, who undertook the RSVP task and also received 1 mA LH stimulation, did not show training transfer. Therefore, it was the specific combination of multitasking training with tDCS that resulted in improved visual search speed. The lack of transfer for the 2mA LH group excludes the role of peripheral^49^ or arousal^50^ effects of stimulation.

Regarding the MVPA data quality, we used an in-scanner motion reduction technique and conservative motion exclusion thresholds, reducing the likelihood of spurious associations which can arise from imaging artefacts. The current findings therefore reinforce the potential effectiveness of tDCS, while also highlighting the dependency of induced effects on the concurrent task being performed, and the intensity of stimulation^22,51,52^. Finally, our neuromodulation interventional study design, using established computerised cognitive measures, enhances the reliability of our findings, and avoids issues related to sample size pervasive in other types of brain-behaviour studies^53^.Previously, the reduction of multitasking costs via extensive training was found to be associated with greater segregation of task-related information processing in predominantly frontal, parietal, and subcortical regions^10^. However, we found no reduction in multitasking cost after training for any of the groups that received tDCS. It may simply be that the training protocol used in the current study - which had 1/3 of the trial numbers used in Garner and Dux^10^ - was not sufficient to induce the appropriate learning for such associations to be observed. However, purely training induced cost improvements were observed for the sham group here. Future work is required to determine whether there are different time scales for task-specific and transfer learning effects in the brain, and the potential disruptive effects of tDCS on task-specific training.

The influence of paired training and tDCS on the visual search task were present for both the left and right hemisphere 1mA groups. Critically, however, only left hemisphere tDCS induced changes in task representations that were associated with improvements in visual search performance. This may reflect dissociable mechanisms for transfer based on the prefrontal circuitry stimulated. The left and right prefrontal cortices have been differentially implicated in many cognitive operations, including motor sequence learning^54^, multitasking^13,55^, planning^56^, and relational integration^57^. Indeed, the integration of relationships between stimuli, or in other words, the abstract rules associated with the task, has been found to be encoded in the left prefrontal cortex^58,59^. Meanwhile, the sharing of these abstract rules has been proposed as a mechanism for transfer^6,60,61^. It may be that the application of tDCS to this region enhances its capacity for sharing and integrating information pertaining to task contingencies, within task-general frontal, parietal, and cerebellar regions, resulting in enhanced generalisation of learning. However, none of the other seven tasks tested demonstrated an effect of tDCS on generalisable improvements in reaction times. It may be that the visual search task was the only task sensitive enough to detect an effect; greater tDCS-specific improvements on the most difficult condition in this task supports this account.

It is also conceivable that tDCS facilitated learning generalisation via improvements in some shared cognitive component of the trained task and visual search. For example, the dorsolateral prefrontal cortex - used as the focal point of stimulation in the current study - is involved in evidence accumulation processes^62^ and the enhancement of information processing after training^8^. Using a linear ballistic accumulator to model response times, Filmer et al^30^ found that dorsolateral prefrontal cortex tDCS induced improvements on both a trained task and visual search, and these changes were related to improvements in information processing efficiency in both tasks. In combination with the current results, these findings suggest that the trained-on multitasking paradigm and the visual search task share central information processing resources, perhaps related to stimulus-response mapping^63^, that can be reduced with combined tDCS and training. Future work is needed to disentangle the specific action of tDCS on the cognitive components associated with performing these tasks.

In conclusion, our findings reveal for the first time that combining prefrontal cortex tDCS with concurrent multitasking training modifies the dynamics of task representations, leading to generalised and lasting improvement on a spatial attention task. These results are consistent with recent theories that emphasise a pivotal role for shared task-representations in knowledge generalisation.

## Materials and Methods

### Study design

The study design was preregistered at the Open Science Framework (https://tinyurl.com/5h8u72j5). The study comprised ten sessions: pre-training, post-training, and follow-up behavioural testing sessions; four sessions combining tDCS with training; and three imaging sessions. The imaging sessions involved initial magnetic resonance spectroscopy, T1-weighted, fluid attenuated inversion recovery, quantitative susceptibility mapping, and diffusion weighted structural scans. We also conducted task-based and resting state functional scans before and after training. In the current study we report exclusively on the task-based functional MRI and T1-weighted structural scans.

The testing sessions (pre-, post-, and follow-up) included an extensive set of nine cognitive tasks aimed at testing different cognitive processes, described below. We conducted these testing sessions on 2014-model Mac mini computers (2.8 GHz Intel Core i5, OSX High Sierra v 10.13.6), with 24-inch ASUS VG248 monitors (144Hz refresh rate) displaying the tasks. We implemented the tasks using MATLAB 2016b (The MathWorks, Inc., Matick, MA) with custom code and the Psychtoolbox^64-66^ (see http://psychtoolbox.org). During the behavioural sessions, participants were positioned approximately 57 cm from the screen.

We used a mixed design, dividing participants evenly among five groups while ensuring their participation in all sessions. Two groups—those receiving sham left prefrontal cortex (PFC) stimulation and those receiving 1 mA left PFC stimulation with single/multi-task training—were double-blinded throughout the experiment. Note that all ‘targets’ for stimulation refer to the anode electrode location. The remaining three groups acted as active controls: a group receiving 2 mA left PFC stimulation with single/multi-task training served as a dosage control; a group receiving 1 mA stimulation to the right PFC with single/multi-task training served as a montage control; and a group receiving 1 mA to the left PFC with rapid serial visual presentation (RSVP) task training served as a task control (active control group). To minimize the impact of task order on baseline performance measurements, we pseudorandomised and counterbalanced task order across all participants and groups.

### Participants

Our sample size and recruitment stopping rule were calculated using a power analysis via G*Power. This analysis was based on conservative effect sizes (Cohen’s D of 0.3) for the behavioural impacts of tDCS on multitasking training, as indicated by our lab’s prior studies^22,30^. This analysis revealed that 33 participants per group would provide 95% power for between-group comparisons, and 46 participants per group would be necessary for the same power for individual differences analyses. Therefore, we set a maximum goal of 250 participants (50 per group) to enable both between- and within-participants analyses.

The COVID-19 pandemic and associated lockdowns resulted in a final sample size of 207 participants. Despite setting a stopping date of September 28th, 2021, we continued data collection until January 15th, 2022, to reach the minimum sample size for between-participants comparisons. This extension was decided without examining the data and was based on the a priori power analysis. After excluding 29 participants for reasons described below, we arrived at a final sample of 178 participants who completed sessions 1 through 9. Of these, 167 participants also completed session 10 (the 1-month follow-up).

### Participant exclusion and group allocation

We excluded six participants due to incidental findings (all pineal gland cysts) in their initial MRI, ten participants who could not tolerate the MRI scans, and three participants who could not complete sessions due to COVID-related lockdowns. Other exclusions included three participants failing to meet the 60% accuracy threshold in the multitasking paradigms at baseline, two participants who missed sessions, and a few others due to personal reasons (1), inappropriate behaviour (1), tDCS discomfort (1, no adverse effects noted), psychoactive medication use before a tDCS session (1), or significantly reduced accuracy in a training session (1, <30% accuracy during pre-training familiarisation). Eight participants couldn’t attend the 1-month follow-up session due to COVID-19 lockdowns, and three didn’t return for testing.

Participants passed a tDCS safety screening questionnaire and a quiz to confirm their understanding of the study’s techniques (i.e., tDCS and MRI) before testing sessions. They also completed an MRI safety questionnaire before MRI sessions. We excluded potential participants before testing based on factors like history of brain trauma, current use of psychoactive medications, and personal or family history of epilepsy. The study was approved by The University of Queensland Human Research Ethics Committee, and all participants provided written informed consent for each session. Participants received compensation at a rate of $20 AUD/h, yielding approximately $380 in total if all 10 sessions were completed. Participants were compensated for their time, regardless of study completion.

After session one, participants were assigned to groups using a custom automated group assignment algorithm. This algorithm aimed to minimise potential baseline group differences by pseudo-randomising participant allocation to groups, based on key characteristics such as age, sex, intervention time (AM or PM), single- and multi-task reaction times, and task completion order.

### Stimulation protocol

Stimulation was delivered using a NeuroConn stimulator, which used two 5 x 5 cm saline-soaked sponges with rubber electrodes secured by rubber straps. For individuals receiving left PFC stimulation, the anodal electrode was positioned 1 cm posterior to F3, and the cathodal electrode was placed over the right supraorbitofrontal cortex, in alignment with prior studies^21,22,67^. Individuals in the right PFC stimulation group had the anodal electrode placed 1 cm posterior to F4, and the cathodal electrode over the left supraorbitofrontal cortex, mirroring the left hemisphere montage. Participants received 13 minutes of online stimulation during training, with a 30-second linear ramp-up and ramp-down. Given previous evidence suggesting that brain stimulation outcomes can be influenced by time of day^68^, participants completed their sessions at roughly the same time each day (±2 h). Our chosen intensities, 1 mA and 2 mA, have been used in most previously published tDCS studies (95%)^69^, thus facilitating comparison of intensity and montage effects with those reported previously. Furthermore, our lab has previously identified distinct effects of stimulation intensity on multitasking performance and decision-making processes^22,70^.

### Training multitask

The key training task involved a sensory-motor response selection task^21,22^, consisting of either a centrally presented coloured circle (red: RGB 237 32 36, dark green: RGB 10 130 65, or dark blue: RGB 44 71 151, subtending approximately 2.7° of visual angle) and/or one of three complex tones (as used in Dux et al^55^). Participants responded by pressing designated keys on a keyboard, with each stimulus mapped to a specific key [A, S, D (left hand) J, K, L (right hand); with the index fingers on D and J keys, respectively]. The task included three different trial types: single-visual (a coloured circle; red, green, or blue), single-auditory (one of three complex tones), or a combined visual-auditory multi-task (one coloured circle and one complex tone). Multi-task trials had both stimuli presented simultaneously (0ms stimulus onset asynchrony), consistent with previous multitasking research^8^. Trials proceeded with a fixation square (0.4° diameter) centrally presented for either 600 or 1000 ms (randomly jittered), followed by a 200 ms stimulus presentation. Visual stimuli appeared centrally, while auditory stimuli played in stereo through Sure SRH440 over-ear headphones. Participants had 2200 ms from stimulus onset to respond. Response mappings were counterbalanced across participants and groups, with half using their left hand for the auditory task and right hand for the visual task, and vice versa for the other half. In the pre-training session, participants practiced both single- and multi-tasks until they achieved a 70% accuracy cut-off for the multi-task condition, minimising exclusion rates due to poor performance. The initial practice comprised 3 blocks: the first two blocks containing 15 trials of each single task (auditory, then visual), and the third block featuring 30 trials with all trial types (single visual, single auditory, and multi-task) presented 10 times each, in random order. If participants failed to reach 70% accuracy, they repeated the third block until the threshold was met. In all sessions (pre-, post-, follow-up & training sessions), participants completed 240 trials, equally divided among the three trial types (80 single-visual, 80 single-auditory, and 80 multi-task) with a 30-second break at the halfway point. Before each subsequent session, participants completed block 3 of the practice session to refresh response mappings (10 trials each: single-visual, single-auditory, and multi-task). The training aimed to increase the speed with which participants could execute single or multiple decisions (indicating more efficient information processing). Each 240-trial session lasted approximately 13 minutes. The control training task group, who trained on a rapid serial visual presentation task (described below), also performed multitasking paradigm during pre-, post-, and follow-up testing sessions.

### Rapid serial visual presentation training task (control)

To evaluate whether any potential advantages of combining stimulation and training were specifically related to multitasking training, an active control group trained on a selective attention control task: a modified Rapid Serial Visual Presentation task (RSVP)^71-73^. Participants were shown a fixation square of 0.4° diameter for a randomly determined duration of either 0.2, 0.3, 0.4, 0.5, or 0.6 seconds. This was followed by a rapid stream of seven distractor numbers, with the aim being to identify a single target letter embedded within the stream (e.g., 4, 2, W, 5, 3, 8, 9, 7). The height of the numerical and alphabetical stimuli was approximately 0.8° of visual angle. Responses were made via keypress on a standard keyboard corresponding to the displayed letter (possible letters presented: ‘A’, ‘B’, ‘C’, ‘D’, ‘E’, ‘F’, ‘G’, ‘H’, ‘J’, ‘K’, ‘M’, ‘N’, ‘P’, ‘R’, ‘S’, ‘T’, ‘W’, ‘Y’, ‘Z’). Letter position was randomly allocated throughout the session between positions 3 and 6 in the serial presentation. The task consisted of 240 trials, divided into three blocks. Each rest period between blocks was dynamically adjusted to ensure task duration (time spent training) matched the single/multi-task that was trained. The presentation duration was dynamically adjusted for each participant to maintain accuracy at approximately 70%. In the pre-training session, each stimulus was initially presented with a 100ms duration, then reduced in 10ms increments if participants accurately responded to five consecutive trials. If two incorrect trials occurred successively, the presentation duration was increased by 10ms per stimulus. The minimum stimulus duration was 10ms. Each subsequent session began with the final presentation duration (i.e., 20ms) of the pre-training session. Presentation duration served as the dependent variable, with shorter durations indicating better performance. The key distinction between this task and the single/multi-response selection training task is that accuracy and presentation duration were the main performance measures, rather than response time. Thus, this task did not require or test for the speed of responses. Furthermore, this task contained forward and backward masking of the target stimulus which limited performance. Each 240-trial session lasted approximately 13 minutes. This task was also used in the pre-, post-, and follow-up testing sessions as a transfer task for the four groups that trained on the multitasking paradigm.

### Visual search

We employed a visual search task (locating a ‘T’ among distractor ‘L’s) as we have observed transfer on this task in two previous experiments that combined tDCS with cognitive training^21,30^. Trials commenced with a fixation dot of 0.25° diameter, lasting for a randomly determined duration of either 0.4 or 0.6 seconds. Subsequently, the fixation cross vanished, and an array of ‘L’ stimuli plus a single target ‘T’ stimulus appeared (total array subtended 17° of visual angle). These stimuli were displayed equidistant from one another in random positions within the search array. Participants needed to find the ‘T’ rotated either 90 degrees or 270 degrees, among randomly rotated—either 90 degrees or 270 degrees—distractor letter ‘L’s. The two stimuli had an approximate linewidth subtending 0.2° and a height of 0.8° of visual angle. Task difficulty was manipulated by varying the set size: either 8, 12, or 16 stimuli, with 80 trials for each set size, totalling 240 trials. Participants had 3,000ms to press the ‘Z’ key with their left index finger if the ‘T’ was rotated by 270 degrees or the ‘M’ key with their right index finger if the ‘T’ was rotated 90 degrees. A target ‘T’ was present on every trial. Practice consisted of 15 trials before each of the pre-, post-, and follow-up testing sessions; feedback was provided for incorrect responses during practice in the form of a brief auditory tone.

### fMRI task

The task performed within the scanner was similar to the response-selection training task paradigm described above. To optimise the task for the MRI environment and enhance blood oxygen level dependent (BOLD) signal detection, we implemented minor modifications. Specifically, participants performed solely the single-task component, presented in blocks of four trials. Within each block, stimuli were randomly presented but were of the same modality. A 12-second rest period separated each block. Participants completed a total of 8 blocks per modality in each run, with 3 runs in total for each session, resulting in 96 trials per modality per session.

### Imaging acquisition

Scans were acquired with a MAGNETOM 7T Plus MRI scanner (Siemens Healthcare, Erlangen, Germany), equipped with a 64-channel receive head coil (Nova Medical, Wilmington, MA, USA). Anatomical T_1_-weighted images were acquired with a magnetisation-prepared 2 rapid acquisition gradient echo (MP2RAGE) sequence for ultra-high-resolution structural images: 0.75mm^2^ isotropic, TR = 4300ms, TE = 3.38ms. High temporal and spatial resolution functional images were acquired using a 1.8mm^2^ isotropic voxel echo planar imaging multi-band fMRI sequence; Tr = 1000ms, TE = 19.4ms, field of view = 192 x 192mm, flip angle = 60°, 57 interleaved slices (1.8mm thick), providing whole-brain coverage. Stimulus presentation was synchronised with volume acquisition. Participants lay supine in the scanner and viewed the visual display via rear projection onto a mirror mounted on the head coil. Other scans not reported on in the current study were also acquired, see supplementary materials in Wards et al^41^ for details.

### Imaging data

Results included in this manuscript come from preprocessing performed using *fMRIPrep* 20.2.3^74^ which is based on *Nipype* 1.6.1^75^. The preprocessing details below were modified from the fMRIPrep html output boilerplate for a typical participant. The full output is included verbatim in the supplementary materials in Wards et al^41^, as per published recommendations for reproducibility^74^. This boilerplate was generated for one participant, so the total number of T1-weighted images and functional runs differed whenever a participant with missing scan/s was processed.

### Anatomical data preprocessing

For full details of anatomical MRI preprocessing, see the supplementary materials in Wards et al^41^. In short, T1-weighted (T1w) images were corrected for intensity non-uniformity, skull-stripped, segmented into cerebrospinal fluid (CSF), white-matter (WM) and gray-matter (GM), and normalised to MNI space. This was all performed within the fMRIPrep pipeline which included the following functional data preprocessing.

### Functional data preprocessing

For full details of functional MRI preprocessing, see the supplementary materials in Wards et al^41^. In short, each of the 6 BOLD runs per session were preprocessed separately for each participant. First, a reference volume and its skull-stripped version were generated. Then a deformation field to correct for susceptibility distortions was estimated, and the BOLD reference scan was co-registered to the T1w reference. Head motion parameters were then estimated, before any spatiotemporal filtering. Slice timing correction was then applied, and the corrected BOLD time-series was resampled onto their original, native space.

## Statistical Analysis

### Behavioural data analysis

We used Bayesian methods to evaluate the strength of evidence in support of or against null and alternative hypotheses concerning task performance in the cognitive battery before and after training. As stated in the pre-registration, we did not apply corrections for multiple comparisons, since the Bayesian approach provides more conservative comparisons than null hypothesis testing procedures, reducing the risk of type 1 errors^40^. We anticipated that some tasks and groups would show no training effect across our control groups and tasks. Hence, we intended to evaluate the strength of evidence for these null effects. Bayes Factors for the alternative hypothesis (BF_10_) exceeding 3.2 indicated meaningful evidence in favour of the alternative hypothesis relative to the null, whereas a BF_10_ less than 0.3125 signified meaningful evidence supporting the null hypothesis over the alternative^39^.

The following procedures were established before conducting our key hypothesis tests. To ensure sufficient trial numbers for analysis and to confirm proper task comprehension and completion, we excluded participants from specific task analyses (Trained Multitask, Transfer Multitask, Dynamic Dual Task, Go No-Go, and Visual Search) if their average accuracy for either the pre- or post-training session fell below 60%. This is in line with our previous study protocols^21,22,70^. Subsequently, at the individual participant level, outlier removal was undertaken by calculating the interquartile range (IQR) for reaction time distributions, multiplying this range by 1.5, and adding this value to the third quartile (Q3 + 1.5 x IQR). Reaction times that exceeded this threshold were excluded for each participant. Reaction times faster than 0.2 seconds were also excluded, as per common practice. We applied the same Q3 + 1.5 x IQR exclusion threshold at the participant level for each task, where a mean reaction time exceeding this threshold for either the pre- or post-training sessions resulted in that participant being excluded from analysis on the respective task. Our preregistration stated that we would use an exclusion threshold > 3 standard deviations for the trained multitask and did not specify an approach for all other tasks. For the sake of consistency with the trial-level approach, we reported data using the IQR outlier analysis for all tasks. To assess the effect of combined training and tDCS on the trained multitasking paradigm, we employed Bayesian t-tests on the performance change (single- and multi-task reaction time, as well as multitasking cost) between the pre- and post-training session and the pre- and approximately 1-month follow-up session. This approach was followed because we had specific hypotheses for the double-blind groups and active controls. For all transfer tasks, key performance metrics were compared between the pre- and post-training sessions, as well as between the pre- and 1-month follow-up sessions, to evaluate the presence of training transfer.

### Univariate fMRI analysis

Custom Matlab (The MathWorks Inc, 2019) batch scripts were used to conduct all univariate analyses, using SPM12 functions and the Marsbar toolbox^76^. The functional MRI data (only for the univariate analysis) were first smoothed using a 6mm Gaussian kernel to reduce spatial noise. We then carried out a first-level within-subject analysis where blood oxygen level dependent (BOLD) signal change was modelled for each single-task performed in the scanner. As visual or auditory tasks were executed within blocks of 4 unimodal trials, these blocks were modelled through a general linear model (GLM) as box-car functions, convolved with a canonical haemodynamic response function (HRF) and its temporal and dispersion derivatives. We also included regressors for six motion parameters (X, Y, Z, pitch, yaw, roll). Each block was modelled as a single trial of 12 seconds duration (4 trials of 3 seconds each). This approach was designed to improve the signal-to-noise ratio for each block. We then conducted a group level statistical parametric map (SPM) analysis on the pre-training and post-training fMRI data separately to identify the regions of interest (ROIs) for subsequent analyses. The coordinates for the maximally activated voxel within a cluster of voxels (10 voxel cluster extent threshold, uncorrected p value = 0.001) that exhibited increased activity for both auditory and visual stimulus blocks, relative to rest periods, were taken to define the ROIs. These ROIs were defined based on the pre-training conjunction map, but using the post-training conjunction map didn’t result in different ROIs.

### Multivariate pattern analysis (MVPA)

To conduct the subsequent MVPA using a linear support vector machine (SVM) on these ROIs separately, as in Garner and Dux^10^, we defined 7 and 9mm radius spheres around the voxel coordinates of the activity peak in each cluster from the conjunction analysis, and a mask was created for each of these sizes for each ROI using the Marsbar toolbox (Brett et al. 2002). Several of the 9mm ROIs overlapped in both cortical (E.g., parietal lobe) and subcortical (E.g., putamen) regions, therefore we focused the analysis on the 7mm ROIs. We observed convergence in the results across these two ROI sizes for a subset of the ROIs. MVPA was implemented using the Pattern Recognition for Neuroimaging Toolbox (PRoNTo version 3)^77^, in MATLAB (The MathWorks Inc 2019). Prior to each multivoxel pattern analysis (MVPA), the data for each voxel in a region of interest (ROI) were z-transformed and mean-centred. This involved subtracting the condition mean for the entire ROI from the response in each individual voxel, which controlled for overall differences in signal amplitude between conditions. We trained a series of binary classifiers to differentiate between patterns of activity linked with the visual and auditory single tasks performed in the scanner. Using a leave-one-run-out cross-validation method, for each iteration one run was held out to evaluate the classifiers’ generalisation performance, and the remaining two runs were used to train the classifier. Decoding accuracy was averaged across each of these cross-validation loops, for each ROI, at each session. Post-minus pre-training decoding accuracy was calculated to obtain the change in decoding for each ROI for each participant after training. To ensure the suitability of the data for subsequent correlational analyses, outliers in the pre-training data for each ROI were removed. Specifically, if a participant’s data for a particular ROI exceeded 3 standard deviations from the whole sample’s mean for that ROI, that participant’s data were removed.

### Functional MRI motion analysis

Along with clear instructions to participants to minimise movement as much as possible, we applied a head motion feedback technique to reduce head movement in the scanner^78^. This involved placing medical tape across participants’ forehead, connected to the head coil, with the goal of providing tactile feedback from movement rather than meaningfully constricting movement. We also employed stringent motion correction and exclusion measures to minimise the influence of motion-related artifacts. Volumes with a framewise displacement (FD) exceeding 0.2mm were removed as outliers due to excessive motion^79^. Runs containing 50% or more motion outlier volumes were excluded^80^. If a participant had two or more runs excluded from a session, their data from that session were removed from subsequent analyses pre-vs post-training. As a result, the data from 11 participants were excluded from analyses for at least one session.

### MVPA – behaviour analysis

Changes in decoding in each of the defined ROIs were separately correlated (using Bayesian correlations, custom python script) with the changes in performance on the relevant behavioural metrics. Using the same criterion as for the behavioural analyses, a BF_10_ > 3.2 was taken to indicate meaningful evidence for the alternative hypothesis – in this case that there was a correlation between the change in decoding in an ROI and the change in performance on a particular behavioural dependent variable – while a BF_10_ < 0.3125 signified evidence for the null hypothesis (no correlation). To assess whether these correlations differed between stimulation groups, we used a Fisher Z transformation of the correlation coefficients and performed a t-test on these values. The resulting p-values were FDR corrected and any remaining comparisons with a value < 0.05 were interpreted as indicating significantly different correlations.

## Acknowledgments

We thank research radiographers, Nicole Atcheson and Aiman Al-Najjar, and research assistants Kali Chidley & Zoie Nott.

## Funding

Australian Research Council DP180101885 (PED, JBM); DP210101977 (PED, HLF, JBM).

Department of Defence (Human Performance Research (HPR) Network Partnership (PED, HLF, JBM).

Australian Government Research Training Program Scholarships (SEE & YW).

National Health and Medical Research Council (Australia) Investigator Grant GNT2010141 (JBM).

European Union’s Horizon 2020 research and innovation program under the Marie Sklodowska-Curie, grant agreement No 796329 (KGG).

Australian Research Council Discovery Early Career Research Award DE190100299 (HLF).

## Author contributions

Conceptualization: YW, SEE, HLF, JBM, PED

Investigation: YW, SEE

Visualization: YW

Supervision: HLF, KGG, JBM, PED

Writing—original draft: YW

Writing—review & editing: SEE, HLF, KGG, JBM, PED

## Data and materials availability

Data files are hosted on The University of Queensland Research Data Management System with access available on request.

## Supplementary Materials

**Table S1.**

| MNI coordinates for selected ROIs |  |  |  |
| --- | --- | --- | --- |
| X | Y | Z | ROI |
| -41 | 37 | 29 | L_DLPFC |
| -59 | 7 | 27 | L_Premotor |
| -19 | -3 | 19 | L_Caudate |
| -45 | -5 | 11 | L_Opercular_cortex |
| -25 | -1 | 11 | L_Putamen |
| -27 | 3 | -5 | L_Putamen2 |
| -1 | 5 | 55 | SMPFC |
| -21 | -25 | 19 | L_Thalamus |
| -27 | -9 | 55 | L_Premotor2 |
| -31 | -19 | -3 | L_Putamen3 |
| -27 | -55 | 65 | L_SupPL |
| -63 | -21 | 27 | L_Ant_Supra |
| -45 | -33 | 43 | L_SupraGyr |
| -57 | -41 | 31 | L_InfPL |
| -19 | -67 | 57 | L_SupPL2 |
| -37 | -49 | -31 | L_Cerebellum_Crus |
| -29 | -67 | -23 | L_Cerebellum_VI |
| -1 | -61 | -21 | Cerebellum_Vermis_VI |
| 1 | -59 | 3 | Cerebellum_Vermis |
| 39 | 39 | 25 | R_DLPFC |
| 39 | 47 | 5 | R_AntPFC |
| 25 | 47 | -15 | R_Orbitofrontal_Cortex |
| 31 | -1 | 61 | R_Premotor |
| 59 | 11 | 25 | R_Pars_Opercularis |
| 43 | -3 | 13 | R_Opercular_Cortex |
| 43 | 7 | -1 | R_Insula |
| 25 | 1 | 9 | R_Putamen |
| 17 | 1 | 17 | R_Caudate |
| 27 | 5 | -5 | R_Putamen2 |
| 63 | -17 | 21 | R_SupraGyr |
| 61 | -35 | 25 | R_SupraGyr2 |
| 61 | -41 | -15 | R_Fusiform |
| 47 | 25 | 5 | R_Pars_Triangularis |
| 49 | -41 | 11 | R_SupraGyr3 |
| 49 | -33 | 45 | R_SupraGyr4 |
| 35 | -45 | 41 | R_IPS |
| 35 | -65 | 45 | R_IPS2 |
| 13 | -69 | 53 | R_Sup_Lat_Occ |
| 29 | -17 | 5 | R_Putamen3 |
| 35 | -45 | -33 | R_Cerebellum_VI |
| 25 | -65 | -23 | R_Cerebellum_VI2 |

